# Conserved dimerization architecture in C-type lectins from virus-vector mosquitoes

**DOI:** 10.64898/2026.02.12.705526

**Authors:** Mattia Bertinelli, Rupesh Balaji Jayachandran, Jack Whitehead, Cédric Leyrat, Annabel v. Clanner, Guido C Paesen, Max Renner

## Abstract

C-type lectins (CTLs) play key roles in innate immunity and microbial carbohydrate recognition. In the disease vector mosquito *Aedes aegypti*, the CTLD-S family comprises 34 soluble CTLs whose members are implicated in flavivirus dissemination and microbial homeostasis, yet their structure and organization remain uncharacterized. Here, we combine X-ray crystallography, small-angle X-ray scattering (SAXS), molecular dynamics, and machine learning–based structure prediction to characterize CTLs in *Aedes aegypti*. We determined the crystal structures of four representative CTLD-S proteins: mosGCTL-1, -3, -6, and -20. All crystals featured mosGCTL proteins in an identical homodimer arrangement, positioning both carbohydrate-binding sites on the same molecular face. Dimerization was confirmed in solution and AlphaFold predictions across the entire CTLD-S family indicated that dimer formation may be a unifying feature of mosquito CTLD-S proteins. For one mosGCTL structure, paucimannose glycans bound at a Ca^2+^-dependent site, demonstrating bi-dentate glycan-binding through one dimer. Finally, machine learning based predictions indicated hundreds of possible CTLD-S heterodimers may be viable, with wide-ranging implications for preferred glycan binding through one dimer. Our findings reveal a conserved dimeric arrangement among mosquito lectins that may underpin carbohydrate recognition relevant to vector-pathogen interactions.

## Introduction

Mosquito-borne viruses, such as dengue virus (DENV), West-Nile virus (WNV), or Japanese encephalitis virus (JEV), are major economic and public health concerns (Pierson & Diamond, 2020; Sukhralia *et al*, 2019). The transmission cycle involves cross-species dissemination between vertebrate hosts and mosquito vectors, requiring viruses to be able to infect and replicate within cells of both organisms (Wu *et al*, 2019). Infected female mosquitoes transmit viruses to the mammalian hosts through a blood-meal. In turn, the mosquitoes acquire an infection by biting a viraemic host and ingesting blood which contains circulating virus. Viruses enter and replicate in cells of the mosquito midgut epithelium and spread to the haemolymph and secondary tissues, including to the salivary glands from where they are transmitted to the mammalian host (Rückert & Ebel, 2018). To maintain this transmission cycle, immunological barriers of both host and vector need to be overcome. Unlike vertebrates, insects lack antibody-based adaptive immunity. Instead, they heavily rely on pattern recognition receptors (PRRs) that recognize pathogen-associated molecular patterns (PAMPs), and utilize innate immune defences to fend off viruses (Cheng *et al*, 2016). Recognition of PAMPs by PRRs can lead to the activation of downstream signalling, eliciting innate immune responses.

C-type lectins (CTLs) are an important class of calcium-dependent PRRs that can recognize carbohydrate structures on the surface of pathogens and shape antimicrobial immunity in both mammals and arthropods (Geijtenbeek & Gringhuis, 2009; Ming *et al*, 2024; Brown *et al*, 2018). For instance, in mammals the soluble CTLs SP-A, SP-D and mannose-binding lectin (MBL) can bind to microbial carbohydrates and act as opsonins, promoting phagocytosis of pathogens and complement activation (Brown *et al*, 2018). However, CTLs can also be exploited by viruses which hijack them to mediate attachment to host cells prior to entry and infection. A well-known example in humans is the C-type lectin domain (CTLD)-containing protein DC-SIGN, which interacts with the surface carbohydrates of, among others, DENV, JEV, and Zika virus (ZIKV), thereby acting as an attachment factor to dendritic and liver cells (Dejnirattisai *et al*, 2011; Liu *et al*, 2017b; Pokidysheva *et al*, 2006; Wang *et al*, 2016; Hamel *et al*, 2015). A family of mosquito CTLs has also been shown to affect infection of the arthropod vectors with viruses (Cheng *et al*, 2010; Li *et al*, 2020; Liu *et al*, 2017a, 2014; Chang *et al*, 2024). In the DENV vector-mosquito *Aedes aegypti*, these proteins were initially designated as “mosGCTL” followed by a number. However, more recently, most of these were grouped as “CTLD-S” proteins: a family of 34 related soluble molecules containing a single CTLD (Adelman & Myles, 2018). The *Aedes aegypti* CTLD-S family member mosGCTL-1 was shown to bind to WNV and promote infection of mosquitoes (Cheng *et al*, 2010). Similar observations were made with mosGCTL-3 and DENV (Liu *et al*, 2014), as well as mosGCTL-7 and JEV (Liu *et al*, 2017a). On the flipside, knockout of mosGCTL-2, which is expressed in the salivary glands of female mosquitoes, was found to lead to an elevation of DENV loads in the vector (Chang *et al*, 2024). Finally, multiple CTLD-S proteins were shown to regulate the composition of the mosquito gut microbiome (Pang *et al*, 2016). These studies suggest that CTLD-S family members can promote or restrict microbial dissemination in the mosquito, likely depending on carbohydrate specificity, downstream signalling, and expression patterns.

While our functional understanding of the complexity of mosquito CTLs is rapidly expanding, we have no insight into their molecular organization. Oligomerization of CTLs is a common occurrence and the spatial arrangement of oligomers can determine recognition patterns of microbial carbohydrates (Drickamer & Taylor, 2015). To better understand architecture and carbohydrate binding of mosquito C-type lectins, we used X-ray crystallography, solution scattering methods, and structure prediction to characterize a representative panel of CTLD-S family members from the vector mosquito *Aedes aegypti*. We determined X-ray structures of mosGCTL-1, mosGCTL-3, mosGCTL-6, and mosGCTL-20, which revealed an identical dimer interface in all crystals. In the case of mosGCTL-20 we were also able to resolve two bound glycans consisting of up to 5 carbohydrate residues. We verified the mosGCTL dimerization behaviour in solution using small-angle X-ray scattering (SAXS) and molecular dynamics (MD) ensemble optimisation. In addition, to assess whether the dimerization interface is shared throughout the whole CTLD-S protein family, we used machine learning based structure prediction to generate predictions of homodimers of all remaining family members. Predictions were consistent with a wider conservation of the observed CTLD-S dimerization mode. This dimeric arrangement is unusual and orients the two glycan-recognition sites towards the same surface, indicating a tandem mode of glycan recognition through one dimer. Finally, predictions of CTLD-S family heterodimers suggested that the dimerization interface is also employed in the formation of heterodimers, indicating the possibility of a large combinatorial space of possible ligand recognition.

## Results

### Crystal structures and solution methods show homodimerization of CTLD-S proteins

The 34 members of the CTLD-S family in *Aedes aegypti* encode N-terminal signal peptides (SP), followed by a single CTLD, with an optional C-terminal extension (**Fig. 1A**) (Adelman & Myles, 2018). To maintain consistency with previously published work, we will refer to CTLD-S proteins by their ‘mosGCTL’ nomenclature. We set out to structurally characterize a representative panel of CTLD-S family members: mosGCTL-1 (AAEL000563), mosGCTL-3 (AAEL000535), mosGCTL-6 (AAEL000283), and mosGCTL-20 (AAEL011408). The degree of homology between the proteins spans a broad range, with amino acid sequence identities ranging from ∼29% to 42% (**Fig. 1B**). mosGCTL-1 has been previously shown to be co-opted by WNV as an attachment factor (Cheng *et al*, 2010), while mosGCTL-6 and mosGCTL-20 have been shown to directly bind glycosylated DENV-2 envelope (E) protein (Liu *et al*, 2014).

**Fig. 1.**
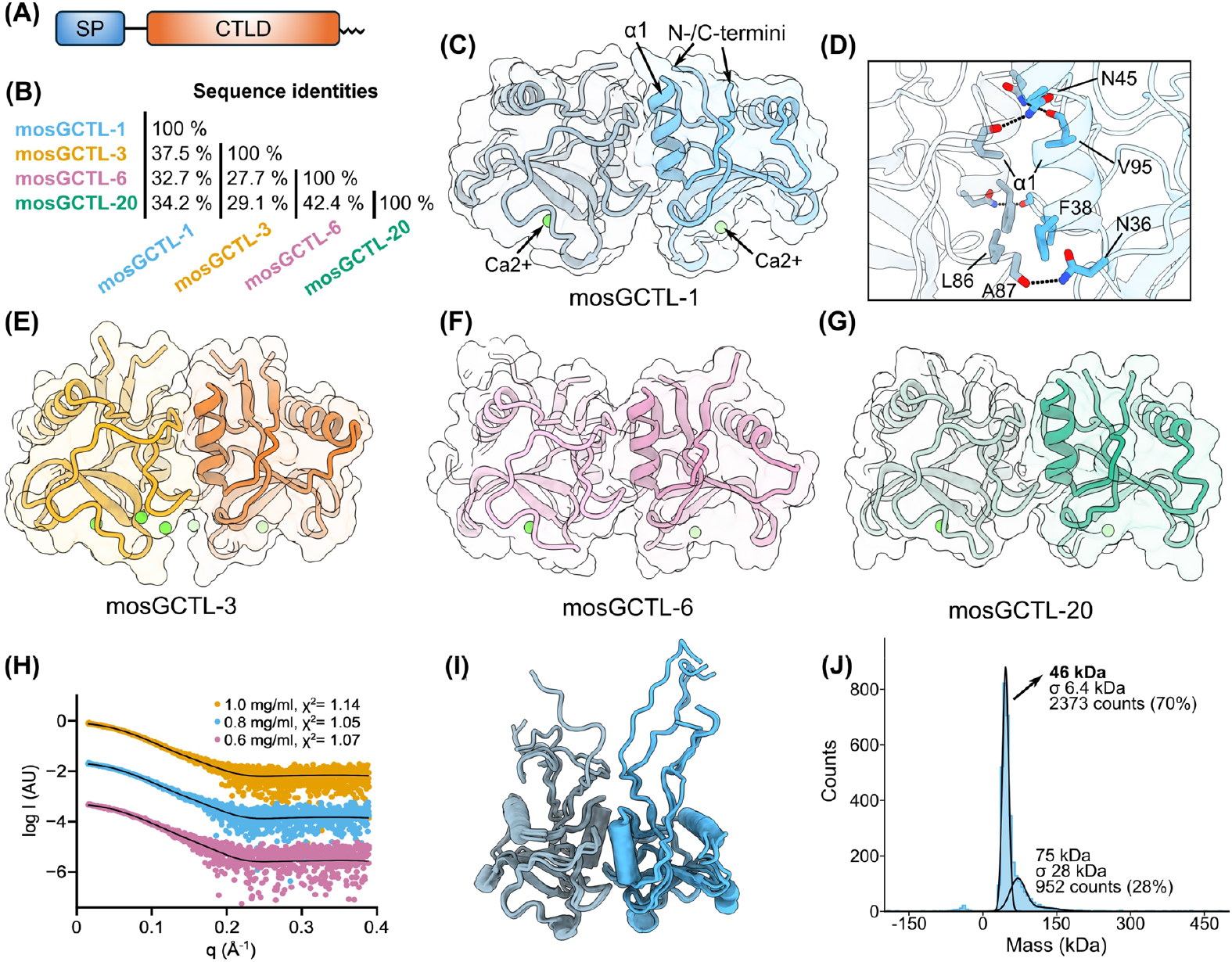
Crystal structures and solution characterization of Aedes aegypti CTLD-S proteins. (A) Schematic organisation of mosGCTLs. The proteins are composed of an N-terminal signal peptide (SP) and a single C-Type Lectin Domain (CTLD). (B) Pairwise sequence identity matrix of mosGCTL proteins that were structurally characterized in this study, highlighting their sequence diversity. (C) Crystallographic dimer of mosGCTL-1. Protomers are depicted in light blue and dark blue. Calcium ions in green. (D) Close-up of the mosGCTL-1 dimer interface. Amino acids forming contacts to the neighbouring protomer are shown as sticks. The helix α1 is indicated. (E-G) Crystal structures of dimers from mosGCTL-3, mosGCTL-6, and mosGCTL-20, respectively. (H) Fitted SAXS curves of mosGCTL-1 at three different concentrations. The fits of optimized ensembles are shown as black lines. (I) Family of models (n = 3) from the optimized ensemble of the 0.8 mg/ml SAXS curve. (J) Mass photometry of recombinant mosGCTL-1 secreted from Sf9 cells. The experiment was performed twice, and a representative plot is shown here.

We solved the crystal structure of mosGCTL-1 at 1.9 Å resolution (**Table S1**) revealing the expected CTLD fold and possessing clear electron density for a bound calcium ion at the carbohydrate binding site (**Fig. S1**). Within the asymmetric unit of the structure, two copies of mosGCTL-1 formed a prominent homodimer (**Fig. 1C**), burying a surface area of ∼1,200 Å^2^. In this configuration the Ca^2+^ atoms of both protomers are positioned at one face of the dimer, while the N- and C-termini are located at the opposite side (**Fig. 1C**). The dimerization interface (**Fig. 1D**) involves the α1 helices of the lectin protomers and is stabilized both by packing of hydrophobic residues (F38 and L86), as well as polar contacts between side-chains and backbone atoms (N36 and A87, N45 and V95). We determined further structures of mosGCTLs-3, -6, and -20 (**Table S1**), and observed the same dominant dimerization interface as in mosGCTL-1 (**Fig. 1E-G**), either within the asymmetric units or via the packing of the crystals. Calculation of the buried surface areas between protomers showed that the extent of the interface area is comparable in all four crystallized CTLD-S proteins (**Fig. S2A**). Inspection of the amino acid composition at the dimerization interfaces via multiple sequence alignment revealed shared interaction patterns (**Fig. S2B**): In all four lectins, polar residues (either N or H) are found at positions 36 and 45 (mosGCTL-1 numbering) and form contacts with backbone oxygens of the neighbouring protomer (**Fig. 1D**). In addition, bulky/hydrophobic side-chains at positions 38 and 86 pack against each other at the dimer interface. The oligomerization mode observed in all four mosGCTLs is unusual and occurs, to the best of our knowledge, only rarely (Brown *et al*, 2007; Huysamen & Brown, 2008).

Next, we sought to probe the oligomeric behaviour in solution. Solution experiments were carried out with mosGCTL-1, as it proved most amenable, with the other mosGCTL proteins showing some time- and concentration-dependent aggregation. As an initial step, we carried out MD simulations of mosGCTL-1 (**Figs S3A** and **S3B**). We then acquired SAXS data of mosGCTL-1, with radii of gyration ranging from 23.5 Å to 24.0 Å, as determined by Guinier fitting (**Fig. S3C**). To confirm the dimeric configuration, we utilized an ensemble optimization method (EOM) approach to fit atomic dimeric models derived from the MD simulation against the experimental SAXS data (Bernadó *et al*, 2007). To this end, 3,000 conformers of the MD simulations were extracted for ensemble optimization, and we obtained good fits of dimeric mosGCTL-1 against the SAXS solution data for all concentrations (**Figs. 1H and S3D**). A sample set of selected conformers (n=3) is shown in **Fig. 1I**. In contrast, it was not possible to obtain good fits to the SAXS curves using only monomeric mosGCTL-1, confirming that the solution scattering data are incompatible with monomeric lectin (**Figs. S3E and S3F**). In addition, we used mass photometry (**Fig. 1J**) to estimate the molecular weight (MW) of recombinant mosGCTL-1 purified from insect cell supernatants (theoretical MW of a monomer: 18.1 kDa). The measured MW (46.0±6.4 kDa) is compatible with a glycosylated dimer. In line with this notion, N-linked glycosylation is predicted at Asn76 of mosGCTL-1 (Gupta & Brunak, 2002). Taken together, our crystal structures and solution data reveal a dimerization architecture that is shared across multiple family members of CTLD-S lectins in *Aedes aegypti*.

### Structure prediction indicates wider conservation of the observed dimerization interface

With experimental evidence of a common dimerization interface for four different mosGCTL proteins, we decided to explore whether there is a wider conservation of this oligomerization mode within the entire family. We used machine learning based structure prediction with AlphaFold 3 (Abramson *et al*, 2024) to predict homodimeric models of the entire CTLD-S family in *Aedes aegypti*. We then plotted the AlphaFold 3 ipTM (interface predicted template modelling) scores of the predicted dimers versus their RMSDs of alignment onto a dimeric crystal structure (**Fig. 2A**). The ipTM score constitutes a predicted accuracy measure of relative positions of proteins within a complex while the alignment RMSD quantifies the structural similarity of the predicted dimers versus an experimental dimer. An example alignment with indicated ipTM and RMSD is shown in **Fig. 2B**. We observed that the majority of predicted dimers cluster at high ipTM scores and that this correlates with high similarity (low RMSD) to the dimer configuration of our crystal structures (green dotted circle, **Fig. 2A**). This suggests that predictions resembling the experimental dimer (**Fig. 2C**) are also more likely to be accurate. In contrast, most dimer predictions with low similarity to the experimental structure also possessed low ipTM scores (red dotted circle, **Fig. 2A**). Inspection of low-ipTM, high-RMSD models showed small interaction surfaces unlikely to be stable in solution (see example in **Fig. 2D**). This may either indicate that the predictions are inaccurate or that for these CTLD-S members oligomerization does not occur. Collectively, AlphaFold 3 predictions of the CTLD-S family in *Aedes aegypti* suggest that the large majority of family members form dimers via the same architecture as observed experimentally through crystallography for mosGCTL-1, -3, -6 and -20.

**Fig. 2.**
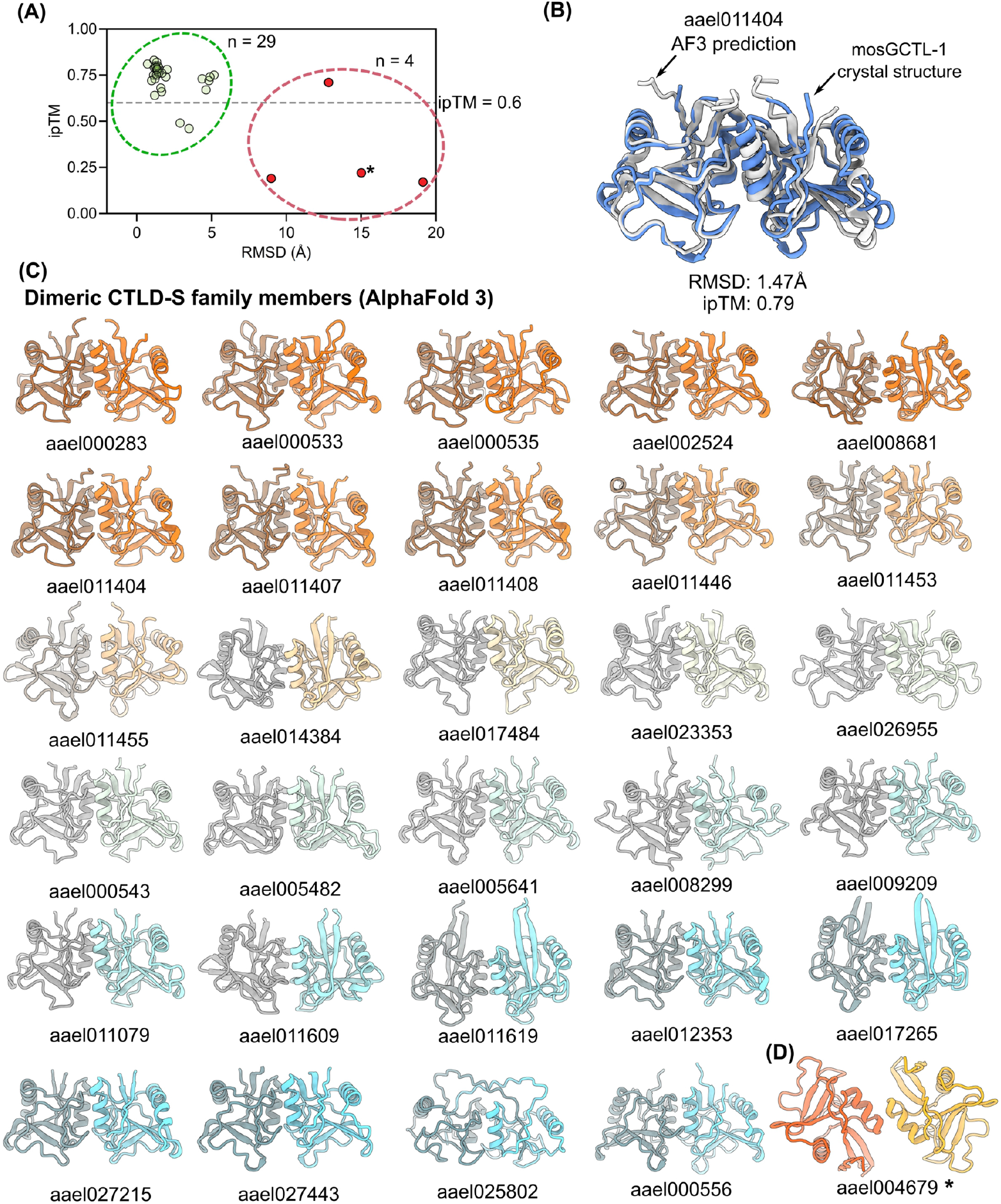
Machine-learning predictions of homodimeric structures of *Aedes aegypti* CTLD-S family members. (A) Plot of prediction ipTM (interface predicted template modeling score) vs. alignment root-mean-square-deviation (RMSD) against the experimental mosGCTL-1 dimer structure. Each data point is the dimer prediction of one CTLD-S member. Explanations of the indicated clusters, see main text. (B) Structural alignment of the prediction of aael011404 (white) against the mosGCTL-1 crystal structure (blue). The comparison highlights the similarity of predicted and experimental dimerization interfaces of different lectins. (C) Tiled view of all structural predictions of 29 dimeric CTLD-S family members within the cluster indicated by green dotted circle in (A). The homodimer predictions indicate a widely conserved dimer arrangement. (D) Representative predicted dimer structure that did not follow the arrangement of the experimental structures and possessed low ipTM (indicated by star in panel (A)).

### mosGCTL proteins possess canonical calcium-binding sites

In C-type lectins, carbohydrates are typically bound by coordinating a Ca^2+^ ion at the ligand binding site via their hydroxyl groups (Drickamer & Taylor, 2015; Zelensky & Gready, 2005). The calcium ion is in turn ligated to the protein via conserved residues at the carbohydrate binding site (Drickamer & Taylor, 2015). The ion thereby serves as an additional coordination bridge between protein and ligand.

In all crystallized mosGCTL proteins (**Fig. 3A and 3B**) we could observe a density compatible with a Ca^2+^ ion coordinated at the long-loop region, in a position referred to as ‘site 2’ in CTLs — the canonical carbohydrate binding site (Zelensky & Gready, 2005). The calcium ion possesses a coordination sphere composed of aspartate/glutamate/asparagine side chains, as well as a backbone carbonyl, occupying six coordinating positions (**Fig 3B**). There are two exceptions, with the site 2 Ca^2+^ of mosGCTL-3 possessing a methionine instead of a glutamate at one position, resulting in only five polar coordinating atoms (arrow in **Fig 3B**). In mosGCTL-20 we also find a serine side chain in the calcium coordination sphere (**Fig 3B**). In addition to the site 2 calcium ion, we could observe two further densities in positions corresponding to site 1 and site 3 in mosGCTL-3 (**Fig. 3C**), although these likely serve structural roles and are not directly involved in carbohydrate binding (Zelensky & Gready, 2005). To further confirm Ca^2+^ binding in solution, we carried out differential scanning fluorimetry (**Fig. 3D**). Supplementation of the buffer with Ca^2+^ ions induced a marked stabilization, shifting the melting temperature of mosGCTL-1 from 74 °C (apo) to up to 87 °C (10 mM CaCl_2_). Conversely, addition of EGTA (a calcium chelating agent) drastically destabilized the protein, reducing the melting temperature to 51 °C (10 mM EGTA). Overall, our crystal structure data of four CTLD-S proteins show the presence of site 2 Ca^2+^ ions, the prerequisite for canonical calcium-dependent carbohydrate binding.

**Fig. 3.**
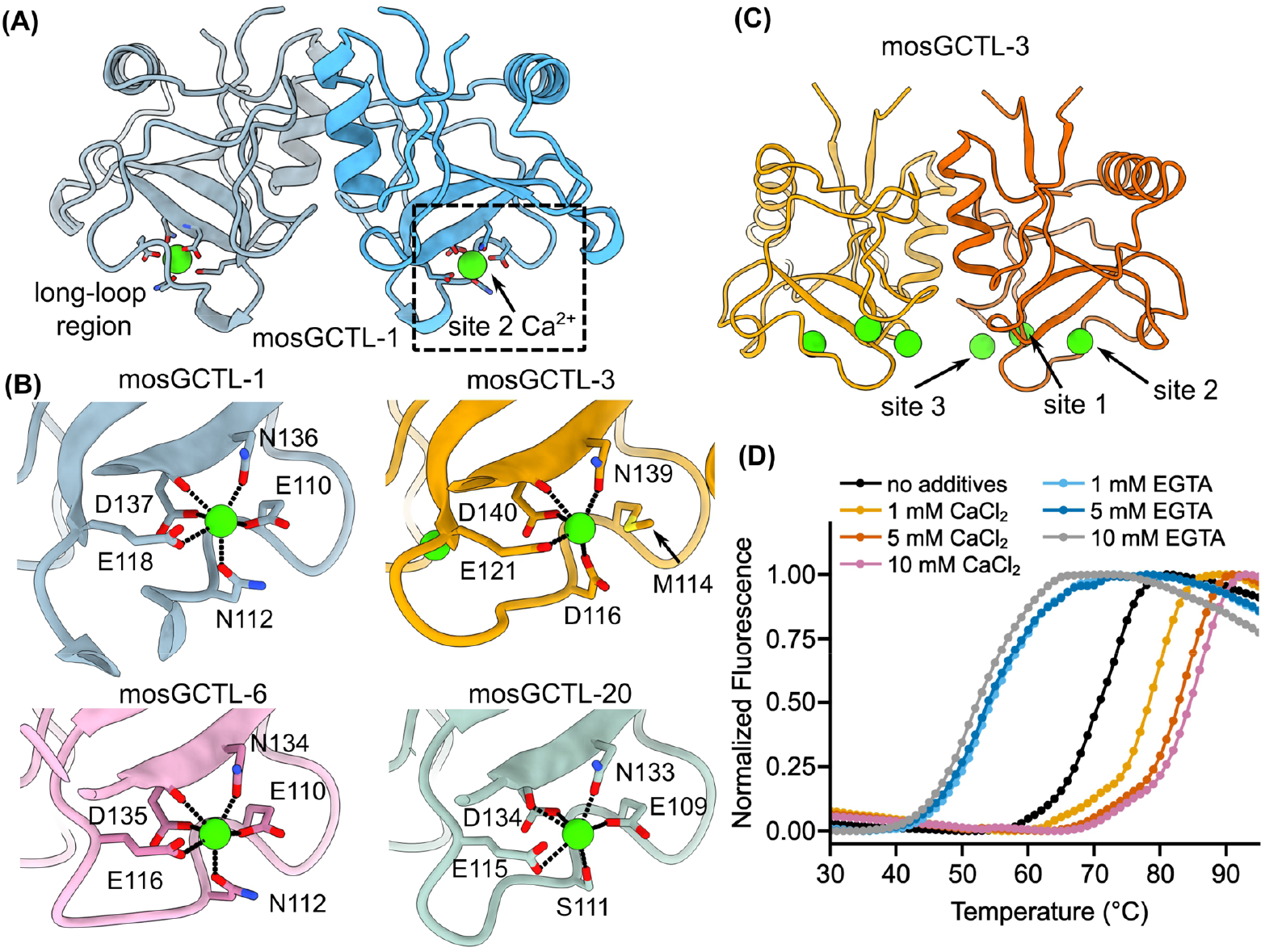
Calcium-binding in mosGCTL proteins. (A) Dimeric mosGCTL-1 structure highlighting Ca^2+^ binding at site 2 at the long-loop region. (B) Close-up views of the site 2 binding sites in mosGCTL-1, -3, -6, and -20, as indicated. Residues coordinating the Ca2+ atom are shown as sticks. (C) Dimeric mosGCTL-3 structure highlighting Ca^2+^ binding at sites 1, 2, and 3. (D) Differential Scanning Fluorimetry curves of mosGCTL-1 in the presence of CaCl_2_ or EGTA at the indicated concentrations.

### Dual glycan binding by the mosGCTL-20 dimer

All *Aedes aegypti* CTLD-S proteins possess N-terminal secretion signals and are thus expected to enter the secretory pathway and potentially undergo glycosylation. We observed the presence of N-linked glycosylation in insect-cell expressed mosGCTL-3 and mosGCTL-20. The glycan of mosGCTL-20, linked to N129, was particularly well-resolved, with good electron density for a full paucimannose structure consisting of two GlcNAc and three Man subunits (**Fig. 4A**). In the mosGCTL-20 crystal structure, we found that the paucimannose tree of one copy of the lectin was bound at the glycan recognition site of a neighbouring copy within the crystal (**Fig. 4B**). This proved serendipitous as it allowed us the determine the structure of mosGCTL-20 in complex with a glycan, without the need of co-crystallization experiments with an externally supplied carbohydrate.

**Fig. 4.**
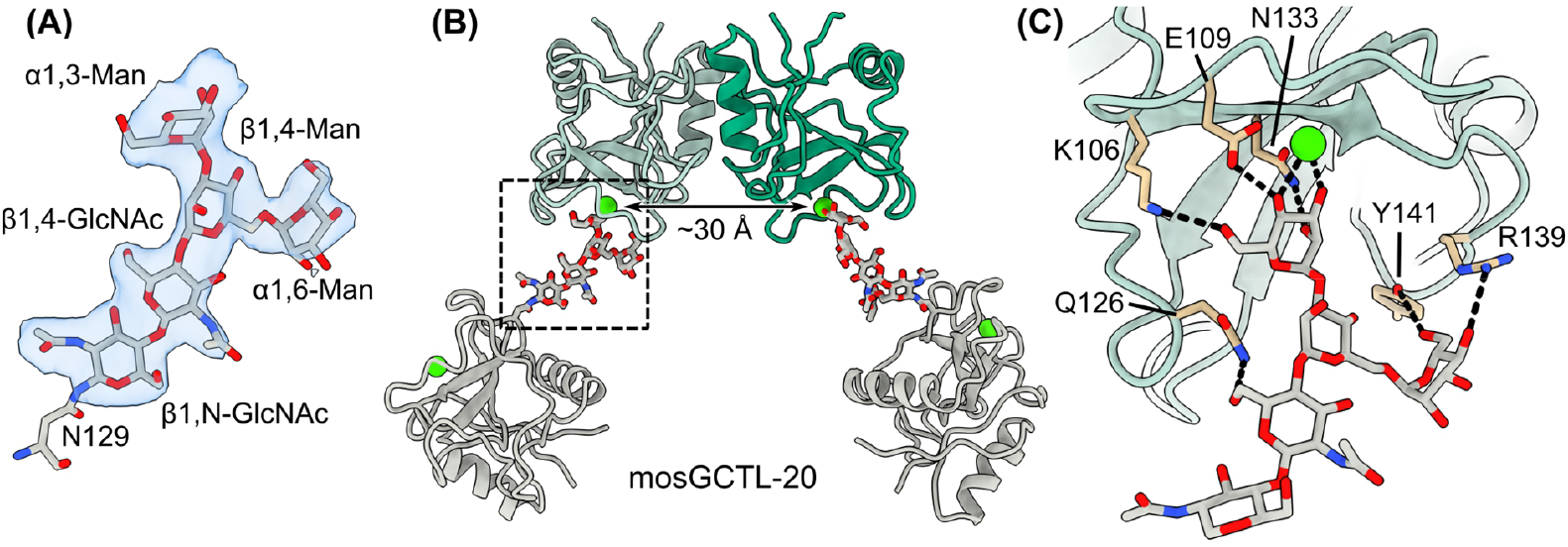
Glycan recognition by mosGCTL-20. (A) Structure and electron density map of a paucimannose glycan resolved in the mosGCTL-20 crystal. The 2Fo-Fc electron density map is shown in blue and contoured at 1.0 σ. (B) Glycan binding within the mosGCTL-20 crystal. Copies of the protein are shown in cartoon representation, glycan is shown as sticks. Ca_2+_ is represented by green spheres. The mosGCTL-20 dimer is shown in shades of green, while the symmetry-related protomers are shown in grey. (C) Close-up of the indicated region from panel (B). Amino acids that contact the paucimannose tree are shown as sticks.

In the context of a dimer, each protomer binds one oligosaccharide originating from separate glycosylation sites (**Fig. 4B**). This places the two glycan recognition sites of a dimer roughly 30 Å apart. The Ca^2+^ atom of mosGCTL-20 is coordinated by the 3- and 4-hydroxyl groups of the terminal α1,3-linked mannose (**Fig. 4C**). The mannose is positioned in such a way that a network of additional polar contacts is formed, involving side chains of K106, E109, and N133 (**Fig. 4C**). Residues surrounding the primary binding site establish additional polar contacts to monosaccharide subunits of the glycan tree. We observe contacts between Y141/R139 and the α1,6-linked mannose (**Fig. 4C**). Finally, Q126 and the 6-hydroxyl group of the β1,4-linked GlcNAc are within hydrogen-bonding distance. Although the observed self-binding of mosGCTL-20 is likely a result of the high local concentrations reached within a crystallization drop, it nonetheless confirms canonical glycan binding at the site 2 Ca^2+^ ion, as well as suggests how one CTLD-S dimer can simultaneously bind two neighboring glycans at around 30 Å distance.

### Heterodimerization of CTLD-S members via a common architecture

While the amount of biochemical data regarding CTLD-S proteins from mosquitoes is limited, one study reported a functional genetic screen that identified two CTLs from the malaria vector *Anopheles gambiae* with an agonistic effect on the development of *Plasmodium* (Osta *et al*, 2004). Follow-up studies showed that the two CTLs, CTL4 and CTLMA2, form a heterodimer and are involved in melanization, an insect innate immune response (Bishnoi *et al*, 2019; Schnitger *et al*, 2009; Nakhleh *et al*, 2017). This prompted us to investigate whether the *Anopheles* CTL4/CTLMA2 heterodimer is formed via the same interface as the *Aedes* mosGCTL proteins. Unfortunately, CTL4/CTLMA2 proved recalcitrant to our efforts of crystallization, however, previous SAXS data has been collected (Bishnoi *et al*, 2019) and is publicly available.

Initially, we utilized AlphaFold3 to predict the structure of the CTL4/CTLMA2 complex. The predicted dimerization mode was remarkably similar to our experimentally determined interfaces for mosGCTL proteins. The CTL4/CTLMA2 complex prediction could be aligned with the mosGCTL-1 homodimeric crystal structure yielding an RMSD of 1.3 Å (**Fig. 5A**). The heterodimer prediction was then used for fitting of the published SAXS data (SASBDB id: SASDFL4). For fitting, we utilized MultiFoXS (Schneidman-Duhovny *et al*, 2010, 2016), allowing conformational sampling of the flexible N- and C-terminal extensions. We obtained an excellent fit (χ^2^ = 1.11) for a single-state model (**Fig. 5B**). The selected model possessed a radius of gyration (Rg = 23.8 Å) that was very similar to what we observed for mosGCTL-1 dimers and showed extended N- and C-termini, again consistent with solution scattering analysis of mosGCTL-1 (compare **Fig. 1I** and **Fig. 5C**). Our analysis indicates that the CTLD-S dimerization mode observed for *Aedes* homodimers may be utilized in other vector mosquito species and in heterodimers.

**Fig. 5.**
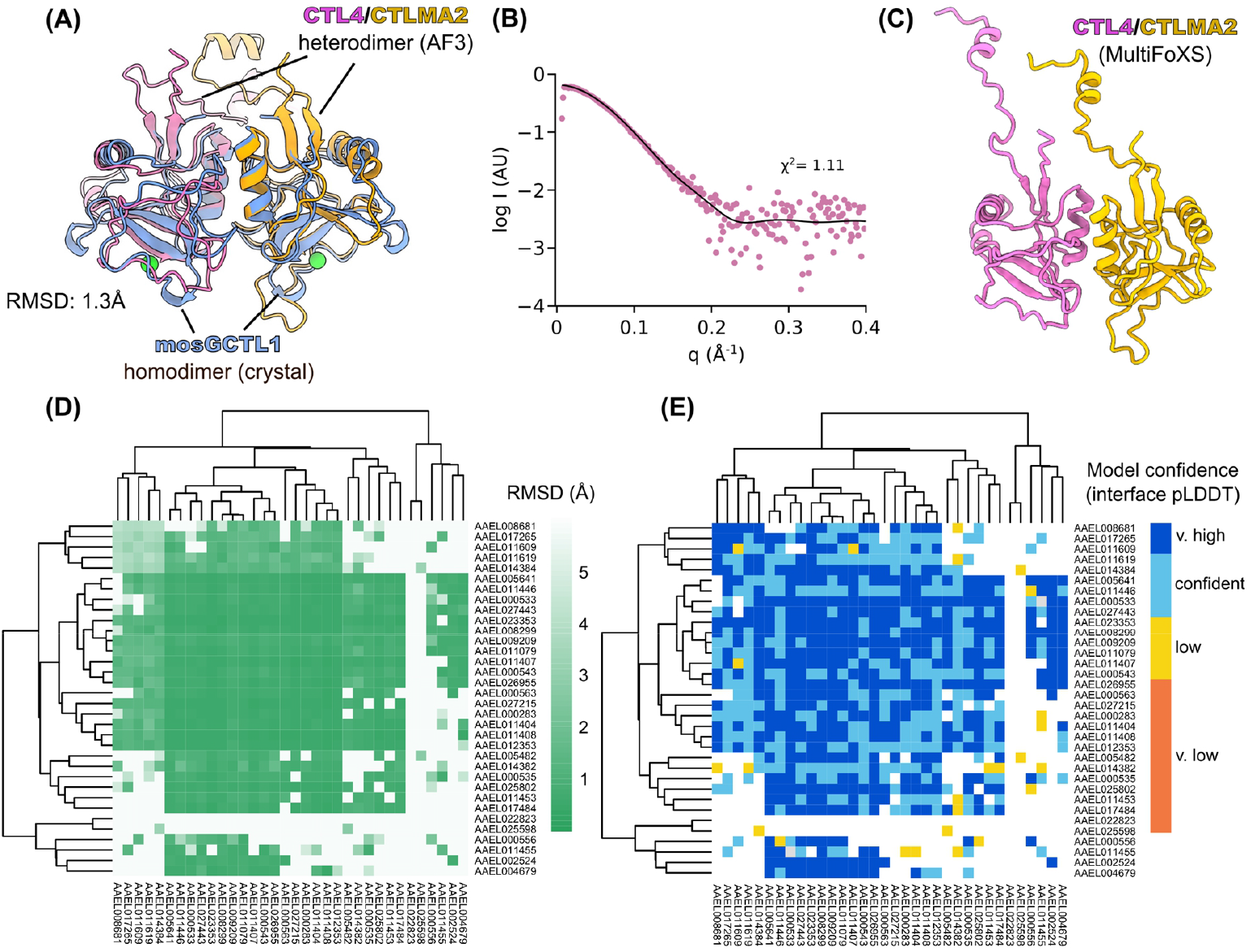
Heterodimerization of CTLD-S proteins. (A) AlphaFold 3 prediction of the CTL4/CTLMA2 complex depicted in pink and yellow, respectively. A structural alignment (RMSD: 1.3 Å) with the mosGCTL-1 homodimer (blue) is shown. (B) MultiFoXS fit (black line) of the predicted CTL4/CTLMA2 heterodimer to SAXS data (SASBD id: sasdfl4, shown in pink). (C) Optimal single-state fit from MultiFoXS corresponding to black curve in (B). (D) Heat maps of RMSD and (E) interface pLDDT scores between *Aedes aegypti* CTLD-S dimer pairs predicted with AlphaPulldown. RMSDs were calculated by structural alignment of predicted vs. experimental dimers and structures with RMSD > 5 Å were filtered out (white boxes). The pLDDT scores shown were calculated from residues at the dimerization interface. Data is hierarchically clustered using Euclidean distance.

Next, we wished to explore whether heterodimerization via this conserved interface may also occur in *Aedes* CTLD-S proteins. We used AlphaPulldown to predict structures of all pairwise combinations of CTLD-S family members (Yu *et al*, 2023). To assess which dimer combinations were predicted with the experimentally observed interface, we compared all structures to the mosGCTL-1 homodimer and calculated RMSD scores, pruning low-RMSD models (white boxes in **Fig. 5D**). In addition, we calculated the average pLDDT score of interface residues to assess the confidence of the dimer predictions (**Fig. 5E**). We identified 392 combinations of possible *Aedes* lectin heterodimers, assembling via a conserved interface, as per our experimental structures. Of these, 380 heterodimer AlphaFold structures possessed interface pLDDT scores indicating ‘confident’ to ‘very high’ model confidence (interface pLDDT > 70, light blue and dark blue in **Fig. 5E**). Our AlphaFold modelling indicates that there may be hundreds of combinations of possible *Aedes* CTLD-S lectin heterodimer pairs which could feasibly assemble.

## Discussion

The ability of CTLs to recognize non-self ligands serves important functions in metazoan immunity and contributes to microbial restriction, however, CTLs may also be subverted as viral attachment factors (Ming *et al*, 2024; Brown *et al*, 2018; Drickamer & Taylor, 2015; Liu *et al*, 2024). In contrast to mammals, arthropods do not possess antibody-mediated adaptive immunity, and so rely heavily on circulating pattern recognition receptors of their innate immune systems (Ming *et al*, 2024). In the *Aedes aegypti* mosquito vector, certain CTL transcripts are induced by flavivirus infection and the encoded proteins have been demonstrated to interact directly with the viral surface glycoproteins (Cheng *et al*, 2010; Li *et al*, 2020; Liu *et al*, 2017a). A number of *Aedes* CTLs have been shown to support dissemination of DENV in mosquitoes while, in contrast, one salivary gland CTL restricted viral loads (Chang *et al*, 2024). In addition, *Aedes* CTL proteins influence bacterial colonization of the mosquito gut (Pang *et al*, 2016). The same study also demonstrated that these proteins can have a protective effect, reducing bacterial elimination by mosquito antimicrobial peptides, under the control of the IMD innate immune pathway. It is clear that there is a complex interplay between arthropod CTL expression patterns, specificity, and microbial restriction and dissemination.

CTLs are typically either secreted or membrane proteins. Commonly, CTLD-containing proteins assemble into certain oligomeric states, and the spatial arrangement of these affects the preference in microbial glycans (Drickamer & Taylor, 2015; Casals *et al*, 2018). For instance, soluble CTLs of the of mammalian innate immunity belonging to the collectin family possess a triple-helical collagen-like domain. The triple helical arrangement leads to higher-order collectin assemblies that typically come in multiples of three: the collectin SP-A assembles into a ‘bouquet’ of six trimers, while SP-D assembles into a cruciform of four trimers.

In this study we investigated the family of 34 related soluble CTLD-S proteins from *Aedes aegypti* using structural, biophysical, and computational approaches. We determined crystal structures of four representative family members spanning a broad range of sequence diversity: mosGCTL-1, -3, -6, and -20. All four structures shared a dominant dimerization interface within the crystals, with key interactions contributed by residues of helix α1. Solution data from SAXS was consistent with the observed dimerization architecture. Furthermore, AlphaFold 3 predictions of all other CTLD-S members showed that the large majority of models form dimers representing our experimentally observed dimer arrangement and that those predictions have the highest confidence scores. Based on these data, we suggest that dimers mediated by helix α1 interactions may be a common organizing principle of *Aedes* innate immune proteins belonging to the CTLD-S group.

To investigate the implications of the dimer architecture in the context of the recognition of viral glycoproteins, we generated a hypothetical model of a mosGCTL dimer bound to the glycans of a flavivirus E dimer — the building block of the virus shell (**Fig. 6**). We found that the spacing and orientation of the lectin protomers were highly compatible with bidentate binding of the N-linked glycans associated with Asn67 of two neighboring E proteins. The arrangement was not dissimilar to the observed dual glycan binding in the mosGCTL-20 crystal structure. However, given the heterogeneity and flexibility glycans can possess (and therefore space they can sample), mosGCTLs may bind to different sets of glycans, possibly across E dimers. This is also consistent with the described heterogeneity of glycan composition in flaviviruses and the observation that DENV serotype 2 can be bound by multiple CTLD-S members (Liu *et al*, 2014; Dubayle *et al*, 2015). In our model, flexible N- and C-terminal tails extending beyond the CTLDs of the lectins are pointing away from the microbial targets (**Fig. 6**) and may very well be involved in the recruitment or activation of downstream effectors such as described with mosPTP-1 (Cheng *et al*, 2010).

**Fig. 6.**
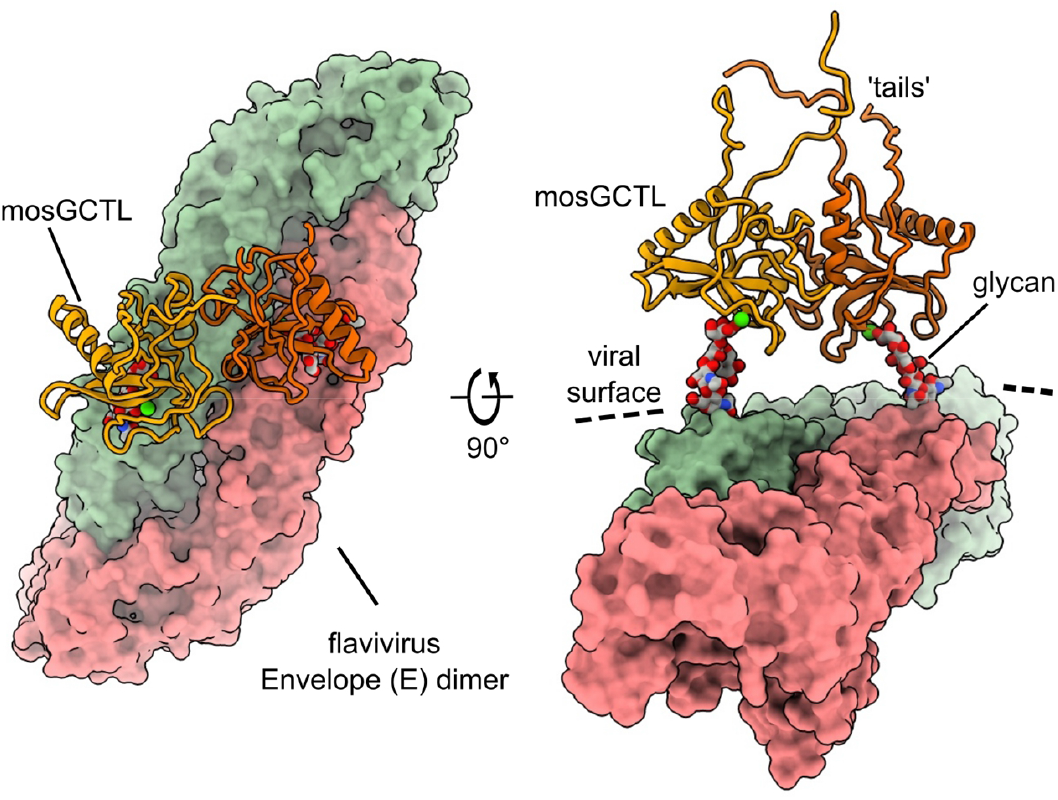
Schematic hypothetical binding mode of a mosGCTL protein (indicated, shown as cartoon) to a flavivirus E glycoprotein (indicated, shown as surface). Glycans are shown as sticks. Calcium as green spheres.

Our AlphaFold 3 predictions in combination with deposited SAXS data suggest that the CTL4/CTLMA2 heterodimer from *Anopheles* uses the same interface for dimerization as in *Aedes* CTLD-S proteins. This prompted us to also calculate structures of all possible dimer pairs in *Aedes aegypti*, resulting in hundreds of predicted viable heterodimer combinations of mosGCTLs (**Fig. 5E**). A previous study indicates that heterodimerization of anopheline CTL4/CTLMA2 changes its ligand specificity as determined by glycan array screening, compared to CTL4 or CTLMA2 individually (Bishnoi *et al*, 2019): CTL4 and CTLMA2 generally bound glycosaminoglycans, however, binding to certain types of carbohydrate ligands (lacto-N-tetraose and lacto-N-neotetraose) could only be detected for the heterodimer. Similar to CTL4/CTLMA2, altered ligand specificity might emerge in mosGCTL heterodimers. The potential formation of hundreds of different heterodimer combinations might lead to a substantial expansion of carbohydrate ligand recognition space in these vector species lacking humoral adaptive immunity.

### Limitations of the study

We have shown evidence of homodimerization of CTLD-S proteins of *Aedes aegypti* in the solution state and in crystals. However, the notion of heterodimerization of these lectins is heavily informed by machine-learning based structure prediction. Further experimental studies will be necessary to confirm whether CTLD-S heterodimerization occurs in mosquitoes, what the possible combinations are, and to elucidate the implications for ligand specificity.

## Materials and Methods

### Cloning, protein expression and purification

Unless noted otherwise, sequences and accession codes of the 34 *Aedes aegypti* CTLD-S proteins were derived from the study of Adelman and Myles (Adelman & Myles, 2018). We expressed *Aedes aegypti* mosGCTL-1, -3, 6, and 20 in Sf9 (*Spodoptera frugiperda*) insect cells. All synthetic genes were purchased from GeneArt (Thermo Fisher Scientific). In the case of mosGCTL-3 the sequence was based on AAEL000535, but included amino acids at the N-terminus, previously identified by 5′-rapid amplification of cDNA ends, which were missing in the entry (Liu *et al*, 2014). Genes of mosGCTL-3, 6, and 20 (excluding the secretion peptide which is included in the vector) were cloned into the pOPING vector (Berrow *et al*, 2007). Ligation-independent cloning was carried out using the In-Fusion system (Takara). The recombinant baculovirus expressing the gene of interest was generated by co-transfecting each plasmid with the flashBACULTRA bacmid (Oxford Expression Systems) into Sf9 cells. Cellfectin II (Thermo Fisher Scientific) was used as the transfection reagent. Recombinant baculovirus was then used to infect Sf9 cells. At 72h post-infection, the culture was harvested (1,000 x g, 10 min, 22 °C) and the supernatant containing the secreted protein of interest recovered and filtered through a 0.22 µm filter. The filtered supernatant was then loaded onto a Ni Sepharose Excel column (GE healthcare) and washed extensively with 25 mM Tris, pH 7.5, 500 mM NaCl. The protein was then eluted from the column using a step-gradient of the same buffer supplemented with 50 mM to 300 mM imidazole. The recovered protein was finally subjected to size exclusion chromatography on a HiLoad Superdex S75 16/60 column (GE Healthcare) equilibrated in 25 mM Tris, pH 7.5, 250 mM NaCl supplemented with 5 mM CaCl_2_.

Additionally, mosGCTL-1 was also procured from GeneArt as a pre-cloned gene insert in a pFastBac1 vector with a pHLsec secretion signal replacing the original at the N-terminus (Aricescu *et al*, 2006), codon optimized for Sf9 cells. Recombinant baculoviruses expressing mosGCTL-1 were produced as recommended by the Bac-to-Bac™ expression system (ThermoFisher Scientific). The pFastBac1 plasmid encoding mosGCTL-1 was used to transform competent *E. coli* DH10Bac™ cells. Recombinant bacmids with the mosGCTL-1 sequence were then purified from successfully transformed cells identified through blue-white screening. The purified bacmids were used to transfect Sf9 cells using ExpiFectamine™ Sf transfection reagent (ThermoFisher Scientific) and the resulting baculoviruses were harvested after 72 hours post transfection. These baculoviruses were passaged twice before being used for recombinant protein expression as above.

Besides the Sf9-based protein production, mosGCTL-1 was also expressed in a bacterial system for crystallography. The gene for mosGCTL-1 (excluding the secretion signal) was cloned into the pOPINE vector (Berrow *et al*, 2007) and expression was carried out in BL21 cells. BL21 cells harboring the expression plasmid were cultured at 37 °C in Luria broth (LB) with appropriate antibiotics, and induced at an OD_600_ of 0.6 by addition of 1 mM isopropyl β-D-1-thiogalactopyranoside for 3 hours. Cells were then harvested (18°C, 20 min, 4000 x g). The pellet was resuspended in 25 mM Tris, pH 7.0, 500 mM NaCl, 5 mM beta-Mercaptoethanol, 8 M Urea and loaded onto a Ni nitrilotriacetic (NTA) agarose (Qiagen, Netherlands) column. After extensive washing, the unfolded protein was eluted in the same buffer, supplemented with 300 mM imidazole. Refolding was carried out by pipetting the sample dropwise into refolding buffer (100 mM Tris, pH 7.0, 1 M arginine, 5 mM reduced glutathione, 0.5 mM oxidized glutathione, 20 mM CaCl_2_, 5 % glycerol), and stirring over night at 4 °C. Then the buffer was exchanged via dialysis to 25 mM citrate, pH 4.0, 500 mM NaCl, 10 mM CaCl_2_. Finally, the refolded protein was subjected to size exclusion chromatography on a HiLoad Superdex S75 16/60 column (GE Healthcare) equilibrated in 25 mM citrate, pH 4.0, 250 mM NaCl supplemented with 10 mM CaCl_2_.

All cloned genes were verified by Sanger sequencing.

### Crystallization, data collection and refinement

Sitting drop vapor diffusion crystallization screens were set up using a Cartesian Technologies dispensing system. mosGCTL-1 crystallized at 298K with the mother liquor condition: 0.2 M proline, 0.1 M HEPES (4-(2-hydroxyethyl)-1-piperazineethanesulfonic acid) pH 7.5, 10 %w/v polyethylene glycol 3350. mosGCTL-3 in 0.2 M magnesium chloride, 0.1 M bis-Tris pH 6.5, 25 %w/v polyethylene glycol 3350. mosGCTL-6 in 0.2 M trimethylamine n-oxide, 0.1 M Tris pH 8.5, 20 %w/v polyethylene glycol monomethyl ether 2000. mosGCTL-20 in 0.5 M sodium succinate pH 7.0, 0.1 M bis-Tris propane pH 7.0. Crystals were harvested and cryoprotected with a solution of the respective mother liquor supplemented with 20% (v/v) glycerol prior to flash cooling in liquid nitrogen. Diffraction data were collected at the I04 and I04-1 beamlines at Diamond Light Source (Didcot, UK) and automatically processed with Xia2 (Winter, 2010). The structure of mosGCTL-1 was solved by molecular replacement using PDB id: 1k9i as a search model in Phaser (McCoy *et al*, 2007). All other mosGCTL proteins were then solved by molecular replacement with mosGCTL-1 as a search model. For each structure, multiple rounds of model building using Coot (Casañal *et al*, 2020) and refinement with Phenix (Liebschner *et al*, 2019) were performed. Statistics for each structure are shown in Table S1.

### Structure prediction

Structures of homodimer lectins were predicted with AlphaFold 3 (Abramson *et al*, 2024), using the online server implementation (https://alphafoldserver.com/). Sequences and accession codes of the 34 *Aedes aegypti* CTLD-S proteins were derived from the study of Adelman and Myles (Adelman & Myles, 2018). For a few cases predictions could be improved by adding Ca^2+^ ions into the system (aael025802, mosGCTL-5, and mosGCTL-23).

For the pairwise prediction of all possible dimer combinations, we carried out an AlphaFold2-based virtual pulldown experiment using AlphaPulldown v0.3 (Yu *et al*, 2023) in all-vs-all mode. We predicted 25 models per pair and used the top-ranking models for further analysis. To quantify the similarity of the predicted dimer architectures vs. the experimentally determined ones, we calculated RMSD values of structural alignments against the crystal structure of mosGCTL-1. We classed predicted dimers with an alignment RMSD of larger than 5 Å as using a different dimerization interface. To assess model confidence of the dimers, we calculated an average pLDDT score (Jumper *et al*, 2021) of the residues located at the interfaces of the respective predictions.

### Differential scanning fluorimetry

Differential scanning fluorimetry experiments were performed in a 96-well format using the Stratagene Mx3005P qPCR System (Agilent Technologies). The SYPRO orange dye (5000x stock; Invitrogen) was used at a final concentration of 3x per sample.

### Small-angle X-ray scattering (SAXS)

SAXS data were obtained at beamline B21 at Diamond Light Source, Didcot, UK. Samples of mosGCTL-1 were concentrated to 1.0 mg/ml, 0.8 mg/ml, and 0.6 mg/ml in 25 mM Citrate, pH 4.0, 1 M NaCl, 10 mM CaCl2. Protein samples and buffer samples were prepared in a 96-well plate and data were collected in batch mode via an Arinax BioSAXS automated sample changer (Grenoble, France). The exposure unit was kept at 15 °C. 120 frames per sample or buffer were recorded at 12.4 keV on a PILATUS 2M detector (Dectris, Baden-Daettwil, Switzerland) with a detector-distance of 4.014 m. A buffer sample was measured after each protein sample. Two-dimensional detector images were processed automatically into one-dimensional SAXS curves via GDA and DAWN (Diamond Light Source, UK). Buffer subtraction, frame averaging, and initial data quality assessment was carried out with ScÅtter, as was the determination of SAXS-derived perimeters.

For the fitting of the CTLMA2/CTL4 heterodimer from *Anopheles gambiae* we made use of previously recorded SAXS data deposited at the Small Angle Scattering Biological Data Bank (SASBDB, https://www.sasbdb.org/) with accession number SASDFL4 (Bishnoi *et al*, 2019). We utilized AlphaFold 3 to predict the structures of the CTLMA2/CTL4 dimer (Abramson *et al*, 2024). Finally, we carried out multi-state fitting of the predicted CTLMA2/CTL4 dimer against the deposited SAXS data, using MultiFoXS (Schneidman-Duhovny *et al*, 2016). We allowed conformational sampling of the N- and C-termini, while defining the dimer of CTLDs as one rigid body, to maintain the oligomerization interface.

### Molecular Dynamics simulations and ensemble optimization

Flexible residues, including the His-tag, at the N- and C-termini of mosGCTL-1 (AAEL000563), which were not visible in the crystal structure, were modelled in an extended conformation with correct stereochemistry. MD simulations were carried out as described previously (Bertinelli *et al*, 2019). Briefly, GROMACS 5 (Abraham *et al*, 2015) was utilized with the AMBER99SB-ILDN* force field (Best & Hummer, 2009; Lindorff-Larsen *et al*, 2010). To prepare the simulation system, mosGCTL-1 was immersed in a box SPC/E water, with a 1.0 nm distance to the edge of the simulation box. Genion was utilized to add 150 mM NaCl to the system. Long-range electrostatics were modelled via the particle-mesh Ewald summation and the P-LINCS algorithm was used to restrain bond-lengths. Virtual sites were used to enable a 5 fs integration time-step. The v-rescale thermostat and the Parrinello–Rahman barostat maintained pressure and temperature at 1 atm and 300 K. Production runs were performed following energy minimization and equilibration of the system. mosGCTL-1 simulations were carried out in triplicate for 100 ns. Average root-mean-square-fluctuations (RMSF) were calculated per-residue within GROMACS. Structural snapshots were extracted from the simulation trajectories every 100 ps, resulting in an overall pool of 3,000 models. A genetic algorithm, implemented in GAJOE (Bernadó *et al*, 2007) was used to select optimized ensembles from the pool of structures, fitting the SAXS data.

### Mass photometry

Prior to mass photometry measurements, affinity-chromatography purified mosGCTL-1 was diluted in a buffer containing 20 nM Tris pH 7.5, 500 mM NaCl, 150 mM Imidazole to a final concentration of 100 nM. Microscope coverslips (No. 1.5H, 24 × 50 mm) were sequentially cleaned with isopropanol and Milli-Q water three times and dried with a clean compressed-air stream. CultureWell™ gaskets (GBL103250-10EA, Sigma-Aldrich) were cut to cover an area of two wells per sample (3 mm diameter, 1 mm deep) and placed in the center of the clean coverslip, ensuring a tight fit through gentle pressure. A droplet of immersion oil (Carl Zeiss™ Immersol™ 518 F, Fisher Scientific) was applied to the objective of the flow-chamber of One^MP^ Mass Photometer (Refeyn Ltd, Oxford, UK), and the prepared coverslip-gasket assembly was mounted and stabilized with small magnets. To each well, 6 µl of fresh dilution buffer was loaded and the “droplet dilution” option was used to determine focus. Then, 14 µl of 100 nM mosGCTL-1 was loaded onto each well, yielding a final concentration of 70 nM for the measurements. Movies lasting 60 s were acquired using AcquireMP (Refeyn Ltd, v2.4.1). All movies were processed and analyzed using DiscoverMP (Refeyn Ltd, v2.4.2). Bovine Serum Albumin was used for mass calibration.

## Acknowledgements

This work was supported by the Wellcome Trust, UK (grants 075491/Z/04 and 204703/Z/16/Z). We are thankful for the generous support by the Kempe Foundation, the Umeå Center for Microbial Research (UCMR), and the Swedish Research Council (grant no: 2025-06548). We thank Diamond Light Source for beamtime and the staff of beamlines I04, I04-1, and B21 for assistance with screening and data collection. We also thank Loic Carrique (University of Oxford, UK), and André Mateus (University of Umeå, Sweden) for critical reading of the manuscript and Farahnaz Ranjbarian (University of Umeå, Sweden) for assistance with mass photometry. Computation used the Oxford Biomedical Research Computing (BMRC) facility, a joint development between the Wellcome Centre for Human Genetics and the Big Data Institute, supported by Health Data Research UK and the NIHR Oxford Biomedical Research Centre. Financial support was provided by a Wellcome Trust Core Award (203141/Z/16/Z). The views expressed are those of the authors and not necessarily those of the NHS, the NIHR or the Department of Health.

## Author Contributions

Conceptualization: MR

Methodology: MB, RBJ, CL, MR

Validation: MB, RBJ, CL, MR

Formal analysis: RBJ, CL, MR

Investigation: MB, RBJ, JW, CL, AvC, GCP, MR

Resources: CL, MR

Writing—original draft: MR

Writing—review & editing: MB, RBJ, JW, CL, AvC, GCP, MR

Visualization: MB, RBJ, MR

Supervision: MR Funding acquisition: MR

## Data availability statement

The atomic coordinates and structure factors reported in this paper have been deposited in the Protein Data Bank (www.pdb.org). The accession numbers are PDBID: 9T9Z (mosGCTL-1), PDBID: 9TA0 (mosGCTL-3), PDBID: 9TA8 (mosGCTL-6), and PDBID: 9TA9 (mosGCTL-20).

## References

Abraham MJ, Murtola T, Schulz R, Páll S, Smith JC, Hess B & Lindahl E (2015) GROMACS: High performance molecular simulations through multi-level parallelism from laptops to supercomputers. SoftwareX 1–2:19–25

Abramson J, Adler J, Dunger J, Evans R, Green T, Pritzel A, Ronneberger O, Willmore L, Ballard AJ, Bambrick J, et al (2024) Accurate structure prediction of biomolecular interactions with AlphaFold 3. Nature 630: 493–500

Adelman Z & Myles K (2018) The C-Type Lectin Domain Gene Family in Aedes aegypti and Their Role in Arbovirus Infection. Viruses 10: 367

Aricescu AR, L. W & Jones EY (2006) A time- and cost-efficient system for high-level protein production in mammalian cells. Acta Crystallographica Section D: Biological Crystallography 62: 1243–1250

Bernadó P, Mylonas E, Petoukhov MV, Blackledge M & Svergun DI (2007) Structural Characterization of Flexible Proteins Using Small-Angle X-ray Scattering. Journal of the American Chemical Society 129

Berrow NS, Alderton D, Sainsbury S, Nettleship J, Assenberg R, Rahman N, Stuart DI & Owens RJ (2007) A versatile ligation-independent cloning method suitable for high-throughput expression screening applications. Nucleic acids research 35: e45

Bertinelli M, Paesen GC, Grimes JM & Renner M (2019) High-resolution crystal structure of arthropod Eiger TNF suggests a mode of receptor engagement and altered surface charge within endosomes. Communications Biology 2

Best RB & Hummer G (2009) Optimized Molecular Dynamics Force Fields Applied to the Helix-Coil Transition of Polypeptides. The Journal of Physical Chemistry B 113: 9004– 9015

Bishnoi R, Sousa GL, Contet A, Day CJ, Hou C-FD, Profitt LA, Singla D, Jennings MP, Valentine AM, Povelones M, et al (2019) Solution structure, glycan specificity and of phenol oxidase inhibitory activity of Anopheles C-type lectins CTL4 and CTLMA2. Scientific Reports 9

Brown GD, Willment JA & Whitehead L (2018) C-type lectins in immunity and homeostasis. Nature Reviews Immunology 18

Brown J, O’Callaghan CA, Marshall ASJ, Gilbert RJC, Siebold C, Gordon S, Brown GD & Jones EY (2007) Structure of the fungal β-glucan-binding immune receptor dectin-1: Implications for function. Protein Science 16

Casals C, Campanero-Rhodes MA, García-Fojeda B & Solís D (2018) The Role of Collectins and Galectins in Lung Innate Immune Defense. Frontiers in Immunology 9

Casañal A, Lohkamp B & Emsley P (2020) Current developments in Coot for macromolecular model building of Electron Cryo-microscopy and Crystallographic Data. Protein Science 29: 1069–1078

Chang Y-C, Liu W-L, Fang P-H, Li J-C, Liu K-L, Huang J-L, Chen H-W, Kao C-F & Chen C-H (2024) Efect of C-type lectin 16 on dengue virus infection in Aedes aegypti salivary glands. PNAS Nexus 3: pgae188

Cheng G, Cox J, Wang P, Krishnan MN, Dai J, Qian F, Anderson JF & Fikrig E (2010) A C-Type Lectin Collaborates with a CD45 Phosphatase Homolog to Facilitate West Nile Virus Infection of Mosquitoes. Cell 142: 714–725

Cheng G, Liu Y, Wang P & Xiao X (2016) Mosquito Defense Strategies against Viral Infection. Trends in Parasitology 32

Dejnirattisai W, Webb AI, Chan V, Jumnainsong A, Davidson A, Mongkolsapaya J & Screaton G (2011) Lectin Switching During Dengue Virus Infection. The Journal of Infectious Diseases 203: 1775–1783

Drickamer K & Taylor ME (2015) Recent insights into structures and functions of C-type lectins in the immune system. Current Opinion in Structural Biology 34

Dubayle J, Vialle S, Schneider D, Pontvianne J, Mantel N, Adam O, Guy B & Talaga P (2015) Site-specific characterization of envelope protein N-glycosylation on Sanofi Pasteur’s tetravalent CYD dengue vaccine. Vaccine 33: 1360–1368

Geijtenbeek TBH & Gringhuis SI (2009) Signalling through C-type lectin receptors: shaping immune responses. Nat Rev Immunol 9: 465–479

Gupta R & Brunak S (2002) Prediction of glycosylation across the human proteome and the correlation to protein function. Pac Symp Biocomput: 310–322

Hamel R, Dejarnac O, Wichit S, Ekchariyawat P, Neyret A, Luplertlop N, Perera-Lecoin M, Surasombatpattana P, Talignani L, Thomas F, et al (2015) Biology of Zika Virus Infection in Human Skin Cells. Journal of Virology 89: 8880–8896

Huysamen C & Brown GD (2008) The fungal pattern recognition receptor, Dectin-1, and the associated cluster of C-type lectin-like receptors. FEMS Microbiology Letters 290

Jumper J, Evans R, Pritzel A, Green T, Figurnov M, Ronneberger O, Tunyasuvunakool K, Bates R, Žídek A, Potapenko A, et al (2021) Highly accurate protein structure prediction with AlphaFold. Nature 596

Li H-H, Cai Y, Li J-C, Su MP, Liu W-L, Cheng L, Chou S-J, Yu G-Y, Wang H-D & Chen C-H (2020) C-Type Lectins Link Immunological and Reproductive Processes in Aedes aegypti. iScience 23

Liebschner D, Afonine PV, Baker ML, Bunkoczi G, Chen VB, Croll TI, Hintze B, Hung LW, Jain S, McCoy AJ, et al (2019) Macromolecular structure determination using X-rays, neutrons and electrons: Recent developments in Phenix. Acta Crystallographica Section D: Structural Biology 75: 861–877

Lindorf-Larsen K, Piana S, Palmo K, Maragakis P, Klepeis JL, Dror RO & Shaw DE (2010) Improved side-chain torsion potentials for the Amber ff99SB protein force field. Proteins: Structure, Function, and Bioinformatics: NA-NA

Liu J, Quan Y, Tong H, Zhu Y, Shi X, Liu Y & Cheng G (2024) Insights into mosquito-borne arbovirus receptors. Cell Insight 3: 100196

Liu K, Qian Y, Jung Y-S, Zhou B, Cao R, Shen T, Shao D, Wei J, Ma Z, Chen P, et al (2017a) mosGCTL-7, a C-Type Lectin Protein, Mediates Japanese Encephalitis Virus Infection in Mosquitoes. Journal of Virology 91

Liu P, Ridilla M, Patel P, Betts L, Gallichotte E, Shahidi L, Thompson NL & Jacobson K (2017b) Beyond attachment: Roles of DC-SIGN in dengue virus infection. Traffic 18: 218–231

Liu Y, Zhang F, Liu J, Xiao X, Zhang S, Qin C, Xiang Y, Wang P & Cheng G (2014) Transmission-Blocking Antibodies against Mosquito C-Type Lectins for Dengue Prevention. PLoS Pathogens 10: e1003931

McCoy AJ, Grosse-Kunstleve RW, Adams PD, Winn MD, Storoni LC & Read RJ (2007) Phaser crystallographic software. Journal of Applied Crystallography 40: 658–674

Ming Z, Chen Z, Tong H, Zhou X, Feng T & Dai J (2024) Immune functions of C-type lectins in medical arthropods. Insect Science 31: 652–662

Nakhleh J, El Moussawi L & Osta MA (2017) The Melanization Response in Insect Immunity. In

Osta MA, Christophides GK & Kafatos FC (2004) Effects of Mosquito Genes on Plasmodium Development. Science 303

Pang X, Xiao X, Liu Y, Zhang R, Liu J, Liu Q, Wang P & Cheng G (2016) Mosquito C-type lectins maintain gut microbiome homeostasis. Nature Microbiology 1

Pierson TC & Diamond MS (2020) The continued threat of emerging flaviviruses. Nature Microbiology 5

Pokidysheva E, Zhang Y, Battisti AJ, Bator-Kelly CM, Chipman PR, Xiao C, Gregorio GG, Hendrickson WA, Kuhn RJ & Rossmann MG (2006) Cryo-EM Reconstruction of Dengue Virus in Complex with the Carbohydrate Recognition Domain of DC-SIGN. Cell 124: 485–493

Rückert C & Ebel GD (2018) How Do Virus–Mosquito Interactions Lead to Viral Emergence? Trends in Parasitology 34: 310–321

Schneidman-Duhovny D, Hammel M & Sali A (2010) FoXS: a web server for rapid computation and fitting of SAXS profiles. Nucleic acids research 38

Schneidman-Duhovny D, Hammel M, Tainer JA & Sali A (2016) FoXS, FoXSDock and MultiFoXS: Single-state and multi-state structural modeling of proteins and their complexes based on SAXS profiles. Nucleic Acids Res 44: W424–W429

Schnitger AKD, Yassine H, Kafatos FC & Osta MA (2009) Two C-type Lectins Cooperate to Defend Anopheles gambiae against Gram-negative Bacteria. Journal of Biological Chemistry 284

Sukhralia S, Verma M, Gopirajan S, Dhanaraj PS, Lal R, Mehla N & Kant CR (2019) From dengue to Zika: the wide spread of mosquito-borne arboviruses. European journal of clinical microbiology & infectious diseases : official publication of the European Society of Clinical Microbiology 38: 3–14

Wang P, Hu K, Luo S, Zhang M, Deng X, Li C, Jin W, Hu B, He S, Li M, et al (2016) DC-SIGN as an attachment factor mediates Japanese encephalitis virus infection of human dendritic cells via interaction with a single high-mannose residue of viral E glycoprotein. Virology 488: 108–119

Winter G (2010) xia2 : an expert system for macromolecular crystallography data reduction. Journal of Applied Crystallography 43: 186–190

Wu P, Yu X, Wang P & Cheng G (2019) Arbovirus lifecycle in mosquito: acquisition, propagation and transmission. Expert Reviews in Molecular Medicine 21

Yu D, Chojnowski G, Rosenthal M & Kosinski J (2023) AlphaPulldown—a python package for protein–protein interaction screens using AlphaFold-Multimer. Bioinformatics 39: btac749

Zelensky AN & Gready JE (2005) The C-type lectin-like domain superfamily. FEBS Journal 272

