## Supporting Information for "Conserved dimerization architecture in C-type lectins from virus-vector mosquitoes"

**Table S1.** Crystallographic data collection and refinement statistics.

| | mosGCTL-1 | mosGCTL-3 | mosGCTL-6 | mosGCTL-20 |
|---|---|---|---|---|
| **Data Collection** | | | | |
| Space group | P 1 21 1 | P 2 2 21 | P 21 21 21 | P 32 2 1 |
| Cell dimensions | | | | |
| *a, b, c* (Å) | 32.75, 102.13, 86.71 | 90.27, 81.09, 85.79 | 32.51, 89.30, 96.34 | 118.33, 118.33, 103.23 |
| *α, β, γ* (°) | 90.0, 92.6, 90.0 | 90.0, 90.0, 90.0 | 90.0, 90.0, 90.0 | 90.0, 90.0, 120.0 |
| Wavelength (Å) | 0.91587 | 0.97950 | 0.97950 | 0.91587 |
| Resolution (Å) | 51.06-1.89 (1.92-1.89) | 36.66-2.38 (2.42-2.38) | 44.67-1.44 (1.46-1.44) | 102.47-2.86 (2.91-2.86) |
| CC (1/2) | 0.997 (0.523) | 0.999 (0.641) | 0.999 (0.308) | 0.998 (0.411) |
| $R_{merge}$ | 0.154 (1.332) | 0.068 (1.651) | 0.171 (2.983) | 0.422 (3.178) |
| $R_{p.i.m.}$ | 0.065 (0.614) | 0.020 (0.512) | 0.066 (0.817) | 0.097 (0.727) |
| $I / \sigma I$ | 8.5 (1.4) | 19.1 (1.4) | 9.2 (0.6) | 6.1 (1.1) |
| Completeness (%) | 99.6 (99.7) | 100.0 (100.0) | 98.8 (96.3) | 100.0 (99.6) |
| Redundancy | 6.6 (5.7) | 12.8 (11.3) | 13.5 (14.0) | 19.6 (19.9) |
| **Refinement** | | | | |
| No. reflections | 45330 (4527) | 25875 (2519) | 51068 (4932) | 19671 (2756) |
| $R_{work}$ / $R_{free}$ | 19.0 / 21.7 | 21.1 / 25.7 | 16.7 / 19.6 | 22.8 / 26.6 |
| No. atoms | | | | |
| Macromolecules | 4056 | 4199 | 2135 | 3039 |
| Ligands | 48 | 93 | 8 | 195 |
| Solvent | 432 | 43 | 336 | 7 |
| B-factors | | | | |
| Macromolecules | 29.5 | 89.9 | 20.6 | 54.6 |
| Ligands | 34.9 | 116.2 | 18.8 | 80.7 |
| Solvent | 38.4 | 66.6 | 33.2 | 36.5 |
| R.m.s deviations | | | | |
| Bond lengths (Å) | 0.003 | 0.011 | 0.003 | 0.007 |
| Bond angles (°) | 0.57s | 1.73 | 0.68 | 1.13 |
| Ramachandran | | | | |
| Favored | 95.93 | 97.02 | 95.63 | 98.10 |
| Allowed | 4.07 | 2.98 | 4.37 | 1.90 |
| Outliers | 0 | 0 | 0 | 0 |

The values in parenthesis refer to the highest resolution shells.

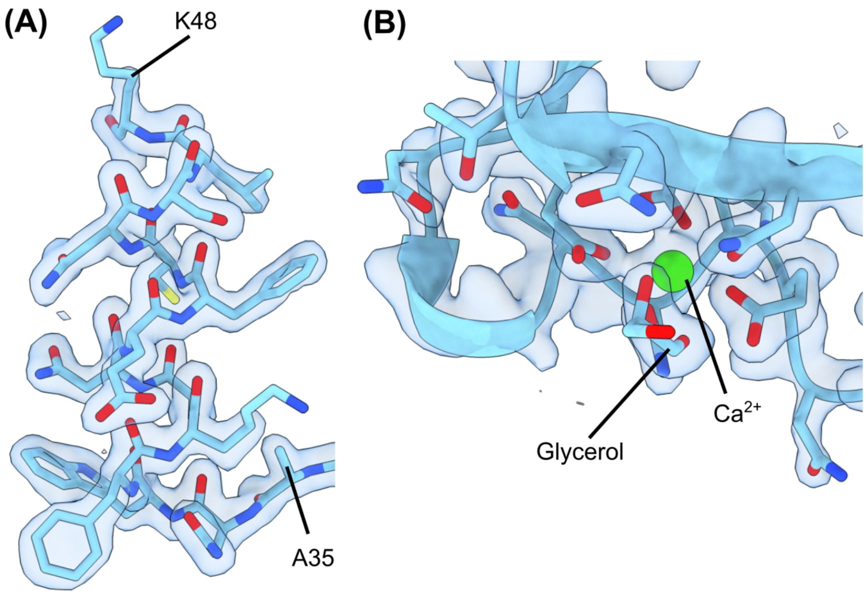


**Fig. S1.** Representative electron density of mosGCTL-1. (A) 2Fo-Fc electron density map is shown in blue and contoured at 1.5 σ for the α1 helix. (B) 2Fo-Fc electron density map is shown in blue and contoured at 1.5 σ for the carbohydrate-binding region.

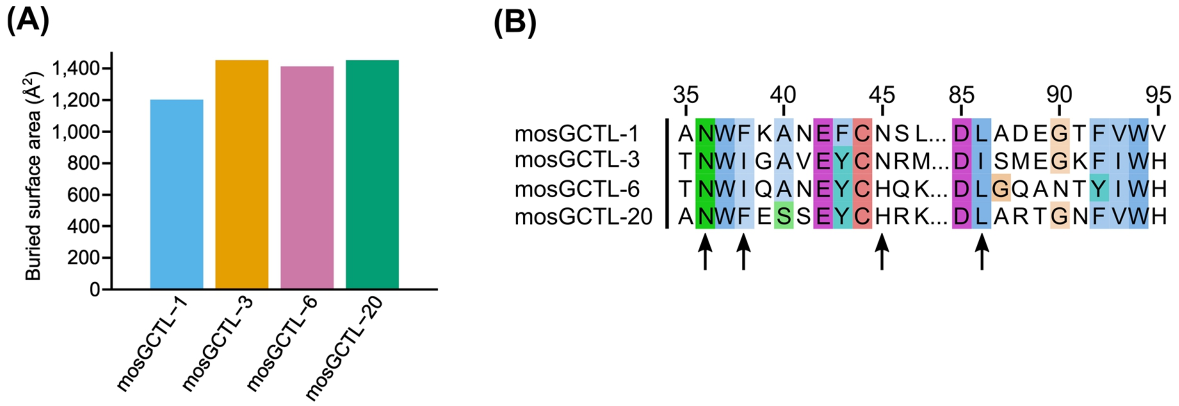


**Fig. S2.** Conservation of the dimer interfaces between experimentally determined mosGCTL structures. (A) Comparison of buried surface areas between mosGCTL dimers calculated from the crystal structures. (B) Multiple sequence alignment of the dimerization regions of solved mosGCTLs. Residues involved in the formation of the dimer interface are indicated by black arrows.

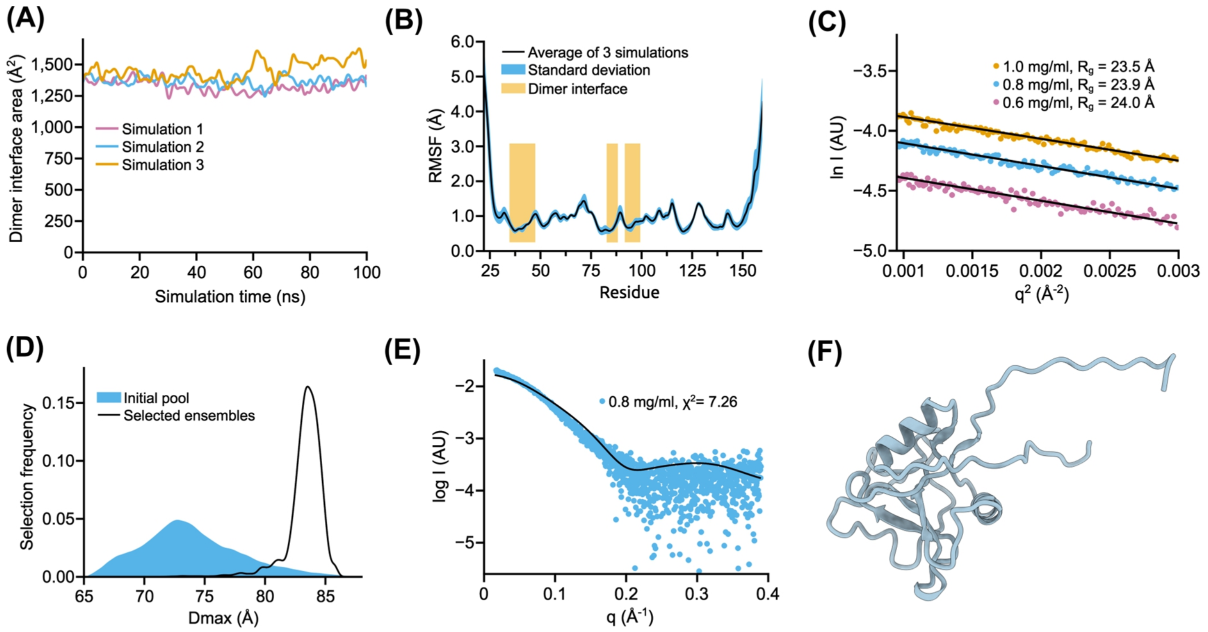


**Fig. S3.** Solution characterization of mosGCTL-1 by MD and SAXS. (A) Development of the dimer interface area over the course of MD trajectories. Three independent replicates are shown. (B) Per-residue root-mean-square fluctuations (RMSF) of Cα atoms averaged over the three MD simulations. The dimer interface regions are indicated. (C) Guinier plots from SAXS data and corresponding fits for mosGCTL-1 at three concentrations. The resulting radii of gyration (Rg) are indicated. (D) Size distributions for the pool of structures generated by MD (blue), used for fitting of SAXS curves, and for the optimized ensembles (black), averaged over all three concentrations. (E) Negative control fitting of monomeric GCTL-1 to the SAXS data resulted in very high $\chi^2$ values, demonstrating that a single protomer cannot account for the solution scattering. (F) Monomeric mosGCTL-1 used for fitting in (E).
